# Diversity and community structure of aerobic anoxygenic phototrophic bacteria are shaped by the deep chlorophyll maximum

**DOI:** 10.64898/2025.12.09.693174

**Authors:** Carlota R. Gazulla, Isabel Ferrera, Vanessa Balagué, Carolina Marín-Vindas, Alba González-Vega, José Escánez-Pérez, Eugenio Fraile-Nuez, Jesús M. Arrieta, Josep M. Gasol, Olga Sánchez

## Abstract

The surface ocean exhibits strong vertical gradients in light, nutrients, and temperature, shaping the phytoplankton distribution which often defines a deep chlorophyll maximum (DCM). Aerobic anoxygenic phototrophic (AAP) bacteria inhabit the euphotic zone, with their abundances following the chlorophyll *a* variability. While AAP bacterial communities are known to differ across regions with contrasting environmental conditions, their vertical distribution remains poorly understood. We hypothesized that the diversity and community structure of AAP bacteria would vary across the vertical gradient, in relation to changes in environmental variables and following the DCM profile. To test this hypothesis, we studied the composition of AAP communities at different depths along the DCM structure in the South and Central Atlantic Ocean, by means of amplicon sequencing of the *pufM* gene. The results show significant differences in richness, community structure, and taxonomic composition of samples from different layers of the DCM, highlighting the dependance of AAP bacteria on its structure. Remarkably, the use of primers with broad phylogenetic coverage enabled the recovery of several phylogroups previously detected only through metagenomics. We show that they represent a significant fraction of marine AAP communities, provide clues on their ecological preferences, and confirm their association with the family *Candidatus* Luxescamonaceae, with genomic potential for carbon fixation.

## Introduction

The stratified pelagic sunlit ocean exhibits pronounced gradients of temperature, salinity, and light irradiance, undergoing relevant changes across tens of meters in the vertical scale. With increasing depth, there is a decline in light intensity and a rise in nutrient concentrations. In stratified oceanic waters, the zone corresponding to the upper nutricline where there is still sufficient light to support photosynthesis harbors the Deep Chlorophyll Maximum (DCM) [1, 2]. The DCM appears as the most distinctive vertical structure that can be observed in stratified waters, situated above the pycnocline and intricately linked to the nutricline [3, 4]. This dynamic layer varies across both space and time [1], and concentrates a substantial fraction of ocean primary productivity [5–7]. Detailed characterization of the fine vertical distribution of phytoplankton groups within the DCM has uncovered its non-uniform structure, revealing the preference of specific phytoplankton groups towards distinct ecological niches [8, 9]. Considering that heterotrophic bacteria rely on phytoplankton-derived organic matter, the structure of the DCM also influence the vertical distribution of different heterotrophic groups. In fact, several studies have compared the microbiota in surface and DCM layers and observed differences in cell abundances, as well as in taxonomic and functional richness [10–12]. However, fine vertical profiling of bacterioplankton along the DCM is limited, with only a handful of studies conducted so far [13, 14, 15].

Besides phototrophs and heterotrophs, marine prokaryotic communities include photoheterotrophs, i.e. proteorhodopsin (PR)-containing bacteria and aerobic anoxygenic phototrophic (AAP) bacteria. These two groups exhibit different distributions in surface waters: PR-containing bacteria are abundant in oligotrophic systems [16, 17, 18] and AAP bacteria thrive better in more eutrophic waters [19–22]. Within the euphotic zone, bacteriochlorophyll *a* (BChl *a*) –the photosynthetic pigment of AAP bacteria– and Chl *a* occupied similar niches along various trophic gradients [17]. In addition, the distribution of AAP cells has been observed to be tightly coupled with Chl *a* and the abundance of cyanobacteria and phototrophic picoeukaryotes [20]. Despite the growing information on the vertical distribution of AAP bacteria, there is little information about the diversity of their communities within the euphotic zone, as most studies have focused on their horizontal variation across the surface ocean [23–28]. These studies show that, in the surface, AAP communities differ across areas with contrasting temperatures, salinity, and chlorophyll levels. We hypothesize that the structure of AAP communities is strongly connected to the DCM profile along the vertical dimension, which displays high environmental variability over a few tens of meters. Notably, the only study examining vertical differences in AAP communities found that populations above the DCM were significantly more similar to each other than to those within or below it [29]. AAP communities are typically composed mainly of Alpha- and Gammaproteobacteria. However, a recent analysis comparing amplicon and metagenomic approaches to study the diversity of the *pufM* gene, the genetic marker of this group, showed that a significant fraction of AAP communities consists on uncultured groups, previously missed in PCR-based studies [30]. Metagenomic analyses have revealed new AAP clades, such as *Candidatus* Luxescamonaceae, which may have the ability to fix carbon [31, 32], and are part of underrepresented uncultured groups distributed across the surface global ocean [30].

In this study, we aimed to assess whether AAP community composition varies along the vertical gradient associated with the DCM. We evaluated the composition of AAP communities by collecting samples at multiple depths across the DCM structure in areas characterized by differing productivity levels in a latitudinal transect spanning the South and Central Atlantic Ocean. The transect encompassed DCM structures of diverse nature, ranging from cold waters with shallow DCMs to warm waters where the DCMs extended below 100 m. AAP communities were investigated through amplicon sequencing of the *pufM* gene using a revised primer set [30]. These primers display a high phylogenetic coverage and minimize primer biases, allowing for a comprehensive representation of AAP communities, including uncultured groups such as *Ca.* Luxescamonaceae, thereby improving our understanding of their ecological distribution patterns.

## Material and Methods

### 1. Sample collection

The Poseidon Expedition, conducted aboard the R/V *Sarmiento de Gamboa* between March and April 2019, spanned approximately 9,000 km along a latitudinal transect (48°S/26°N) in the Atlantic Ocean. Seawater was systematically collected at various depths in 27 stations to capture the structure of the DCM, which was delineated using a CTD profiler SBE 911plus, with samples taken at different depths: surface, above DCM, DCM (defined as the maximal fluorescence point), below DCM, and beyond the end of the fluorescence signal. Additionally, at six stations samples were collected at a higher resolution throughout the DCM (10 sampling depths). At two stations (Station 10 and Station 16), the DCM showed two chlorophyll maxima, and samples were collected at each chlorophyll peak and in between. All samples underwent prefiltration through a 200 µm mesh to remove large plankton. Flow cytometry was used to assess the abundance of cyanobacteria, photosynthetic pico- and nanoeukaryotes, and heterotrophic prokaryotes with high and low nucleic acid content (HNA and LNA), as described in Gasol and Morán [33]. The concentration of inorganic nutrients, specifically nitrate, nitrite, phosphate, and silicate were determined following the procedures outlined in González-Vega et al. [34]. Chlorophyll *a* (Chl *a*) and bacteriochlorophyll *a* (BChl *a*) concentrations were measured using high-performance liquid chromatography and AAP cell abundances were estimated with epifluorescence microscopy (for details, see Gazulla et al. [20]). The mixed layer depth (MLD) was calculated from the profile of temperature differences with depth, using a threshold set at 0.3 °C by every 5 m depth increase (0.06 °C/m). The deepest depth at which the temperature difference exceeded this threshold was identified as the base of the mixed layer.

### 2. DNA extraction, amplification and sequencing of the *pufM* gene

Approximately 2 L of prefiltered seawater were sequentially filtered through 20-μm and 3-μm prefilters onto a 0.2-μm 47 mm polycarbonate filter. Samples were subsequently stored at - 80°C until further processing. DNA was extracted from the 0.2 µm filter (0.2–3 µm fraction) using the phenol–chloroform protocol described in Massana et al. [35]. Partial amplification of the *pufM* gene (∼180 bp fragment) was carried out in 12.5 µL reactions using primers pufM_uniF (GGNAAYYTNTWYTAYAAYCCNTTYCA) and pufM_UniR (YCCATNGTCCANCKCCARAA), from Yutin et al. [36]. The amplification followed the conditions detailed in Gazulla et al. [30]. In brief, PCR these conditions were: an initial denaturation step at 95 °C for 5 min, 35 cycles at 95 °C (30s), 48°C (45s), 72 °C (45s), and a final elongation step at 72 °C for 7 min. DNA sequencing was conducted on an Illumina MiSeq sequencer by AllGenetics & Biology SL (www.allgenetics.eu). Noteworthy, certain samples, particularly some from the DCM and below the DCM, could not be successfully amplified, likely due to the low abundance of AAP bacteria [20]. As a result, we were able to obtain high-quality sequencing results from 84 samples.

### 3. Amplicon sequence variants generation and taxonomic assignation

Primers were removed with cutadapt v3.4 [37] and DADA2 v1.26 [38] was used to discern amplicon sequence variants (ASVs) and eliminate chimeras. To infer the phylogeny of these amplicon sequence variants, we utilized phylogenetic placement with the Evolutionary Placement Algorithm v0.3.5 [39] and a custom made *pufM* database that included sequences from various marine datasets from single-amplified genomes and isolates. The phylogenetic tree was constructed with RAxML-ng [40] and visualized using iTOL [41]. We defined a total of 13 different taxonomic groups based in the initial classification proposed by Yutin et al. [42] and encompassing different orders from the Gamma- and Alpha-proteobacteria. Sequences that could not be further assigned were collectively grouped as “Others”.

### 4. Data analyses

All analyses were conducted in R v4.3.3 (R Core Team 2024). The resulting ASV table underwent rarefaction down to 6,576 reads using the *vegan* package [43]. Diversity indexes (Chao1 and evenness) were estimated using the *vegan* package. To study differences across the vertical and horizontal scale, samples were categorized according to their position along the DCM profile and their corresponding Longhurst provinces [44]. Differences between groups were assessed by analyses of variance with the aov() function in the R Stats Package *stats*, (version 3.6.2) and the post-hoc Tukey’s “Honest Significant Difference” method with function TukeyHDS() from the same package. Correlation between changes in various taxonomic groups and environmental variables were estimated with the rcorr() function (*Hmisc* package, [45]), applying a Bonferroni correction. Relative abundances of taxonomic groups were transformed using the centered log-ratio (CLR) transformation to account for the compositional nature of the data. Pearson correlations between groups were then computed on the CLR-transformed values. For this analysis we included only taxonomic groups with a mean relative abundance above 3%. Ordination tests were based on nonmetric multidimensional scaling (NMDS) analysis using Bray-Curtis (BC) distances, and calculated with the vegdist() function (*vegan* package). The envfit() function (*vegan* package) was used to fit environmental vectors onto the ordination space as well as to test for significant clustering based on Longhurst provinces or the position along the DCM profile. Computing analyses were run at the MARBITS bioinformatics platform at the Institut de Ciències del Mar and at the Picasso Supercomputer at the Supercomputing and Bioinformatics Center of the University of Málaga. The code of the analyses performed can be found at https://gitlab.com/crgazulla/aaps_atlantic_ocean. Amplicon sequences have been deposited in the NCBI Sequence Read Archive (SRA) under BioProject PRJNA1049819.

## Results and Discussion

### 1) Oceanographic context

Most studies targeting the diversity of AAP bacteria have focused on surface ocean samples [24–26, 28, 42, 46] despite evidence that these organisms are distributed throughout the whole euphotic zone [19, 22, 46–48]. These studies, based on AAP cell counts or Bchla *a* measurements, reported highest abundances in eutrophic areas, with strong correlation to chlorophyll distributions. Using the same set of samples than in this study, we showed that AAP bacterial distribution was strongly linked to picophytoplankton (i.e., cyanobacteria and picoeukaryotes) and closely followed the profile of the deep chlorophyll maximum (DCM), with peak abundances typically occurring at or just above the DCM [20]. Noteworthy, AAP cells and BChl *a* were detected from the surface down to depths exceeding 200 m, indicating that there are active AAP communities across the entire euphotic zone.

The Poseidon Expedition went from south to north across four Longhurst oceanographic provinces [44] (Fig. 1A) in the South and Central Atlantic Ocean. It started in an area strongly influenced by the Southern Ocean; Station 2, located in the South Subtropical Convergence Province (SSTC), was characterized by low temperatures and salinity, with the DCM located at approximately 12 m deep (Fig. 1B). Moving northward, the transect entered the South Atlantic oligotrophic gyre (South Atlantic Gyre province, SATL), where both temperature and salinity increased (Stations 3–14). This area was characterized by low surface Chl *a*, deeper DCMs, and low nutrient concentrations. Further north (Western Tropical Atlantic Province, WTRA), closer to the equator (Stations 16–19), DCMs became progressively shallower, with increasing Chl *a* and nutrient concentrations. Beyond the equator, temperatures remained stable while salinity increased as the transect approached the North Atlantic oligotrophic gyre (North Atlantic Tropical Gyre, NATR). This final section was characterized by a strong vertical mixing due to upwelling, and average mixed layer depth (MLD) descending up to 130 m deep (Stations 22–27, mean 101.71 m ± 21.15 m, Fig. S1). More details on the oceanographic conditions can be found in Gazulla et al. [20].

**Figure 1.**
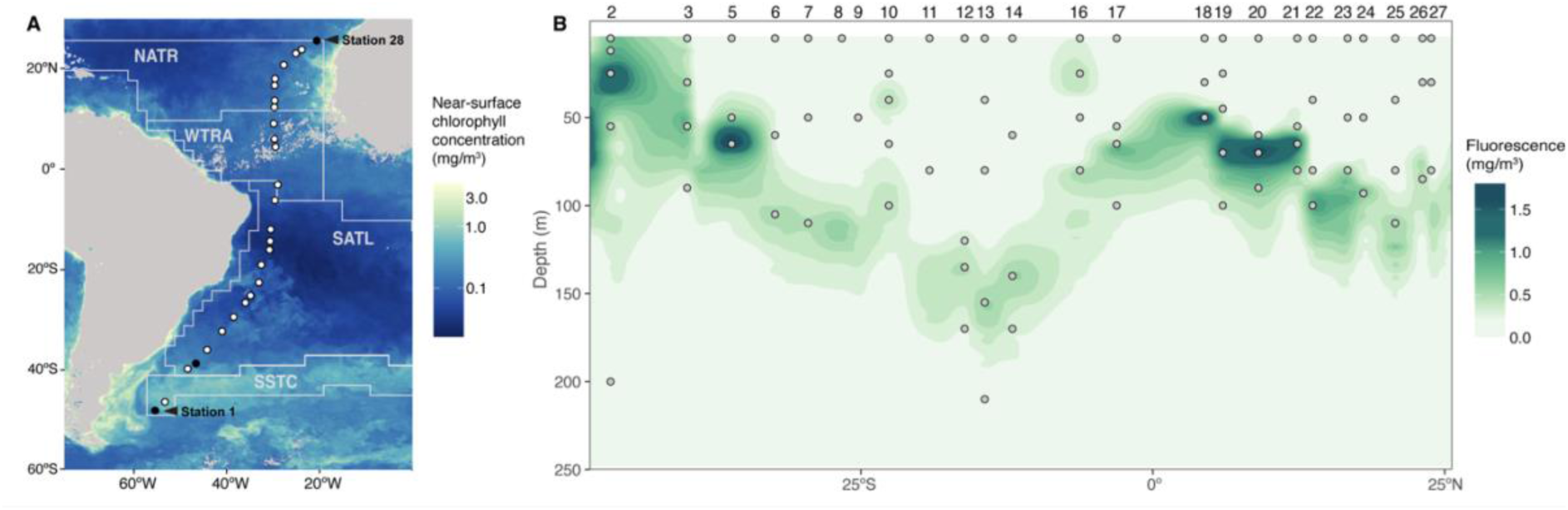
**A)** Near-surface chlorophyll *a* concentrations and station positions along the Poseidon Expedition transect in the South and Central Atlantic (March 2019). Colors indicate chlorophyll *a* (mg·m⁻³) from SNPP-VIIRS data (https://oceancolor.gsfc.nasa.gov/l3/. The transect covers four Longhurst provinces: SSTC, South Subtropical Convergence zone; SATL, South Atlantic gyre, WTRA, Western tropical Atlantic, and NATR, North Atlantic Tropical gyre. **B)** Sectional distribution of *in situ* chlorophyll fluorescence measured by the CTD along the transect. Numbers above the panel indicate station number. Dots indicate the sampling points. Note that station 15 does not exist. Figure adapted from Gazulla et al. [20].

### 2) AAP bacterial diversity and community structure are shaped following the deep chlorophyll maximum profile

We identified a total of 3,350 Amplicon Sequence Variants (ASVs) of the *pufM* gene in the 84 analyzed samples. Chao1 richness values varied between 103 and 339 (mean 214 ± 58), a fair representation of the diversity, as seen by the rarefaction curves (Fig. S2), yet surpassing observations in the same area during the Malaspina Expedition [28] or those in other oceanic sites (e.g., [26, 27, 49]). The high richness values could be explained by the use of primers with broader phylogenetic coverage than previously used ones, as well as by the inclusion of samples from deeper layers of the euphotic zone. In this regard, a total of 1,676 ASVs (49%) were exclusively present below the surface. Previous studies focused exclusively on the surface, ignoring the ocean vertical dimension, and thus missing a high fraction of AAP diversity. Our data show that in most stations, richness reached its maximum where Chl *a* concentration was highest (Fig. 2A, Pearson correlation of Chl *a* and Chao1; n=79, Spearman R=0.39, p<0.001). Interestingly, the two stations with double DCM (Station 10 and Station 16), had different patterns (Fig. S3) and richness reached its maximum at the shallow DCM (45 m) in Station 10 and at the deeper DCM (80 m) in Station 16. Previous studies have also shown that bacterioplankton diversity increases from the surface towards deeper waters in the epipelagic [14, 15]. However it seems that, rather than a gradient of richness with depth, richness of AAP communities follows the distribution of Chl *a,* regardless of the DCM structure. This trend contrasts with previous studies that reported highest diversity values in areas with low Chl *a* concentrations [28, 46] and low inorganic nutrient concentrations [29, 50]. In this study, evenness values were generally high (mean 0.84 ± 0.04, Fig. 2A) and similar across the vertical and latitudinal gradient, being only remarkably low in samples from Station 2 in the SSTC province (Tukey test, p<0.0005). Upwelling processes at the equator also promoted assemblages with high diversity (n=84, Pearson correlation between MLD and Chao1, R=0.39, p<0.001, Pearson corr. between MLD and evenness, R=0.30, p<0.05; Fig. S4AB), and homogenized the composition of the AAP communities across the DCM profile, as seen by the low BC values in stations with a MLD deeper than 100m (Fig. S4C). Finally, the decline of diversity that we observed with latitude aligns well with the global diversity trends observed previously for the whole prokaryotic plankton, where Shannon sharply decreased at high latitudes (above 40°), mainly driven by decreasing ocean temperatures [51].

**Figure 2.**
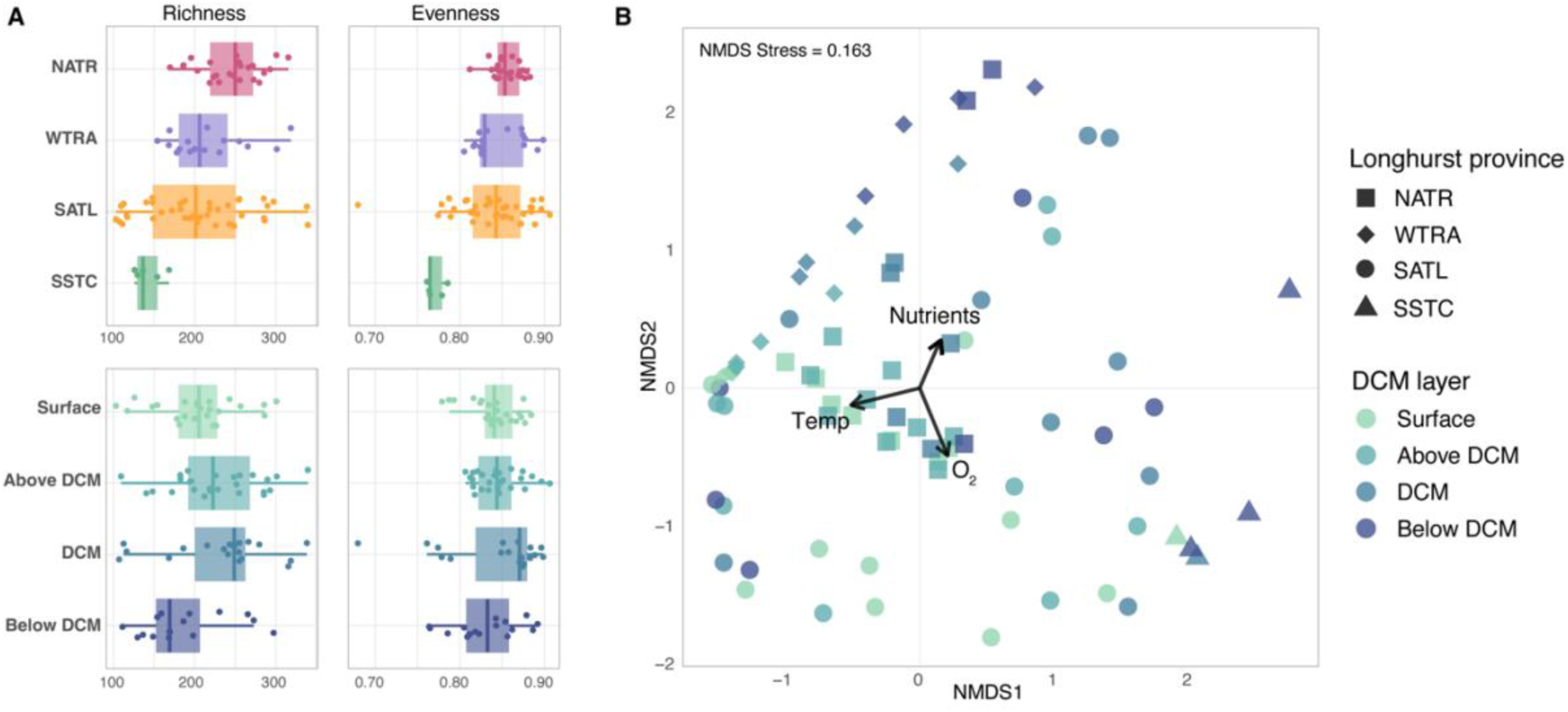
**A)** Alpha diversity measured across latitude (horizontal scale, top panels) and vertically through the Deep Chlorophyll Maximum (DCM) structure (bottom panels). Richness was measured using the Chao1 index. **B)** Non-metrical multidimensional (nMDS) plot based on the Bray-Curtis dissimilarities of AAP community composition. Samples from the distinct DCM layers and Longhurst provinces are denoted by different colors and shapes, respectively. The *envfit* analysis identified temperature (Temp), oxygen (O_2_) and nutrients as the three variables that explained the largest fraction of community variance. SSTC, South Subtropical Convergence zone; SATL, South Atlantic gyre, WTRA, Western tropical Atlantic, and NATR, North Atlantic Tropical gyre.

To explore the biogeographical patterns of AAP communities across the vertical and latitudinal dimensions, we computed a non-metric multidimensional scaling (NMDS) coupled with an *envfit* analysis, to discern the environmental variables influencing the distribution of assemblages (Fig. 2B). Oxygen (R^2^=0.53, p=0.001), temperature (R^2^=0.53, p=0.001) and nutrients (NO_3_^-^, R^2^=0.38, p=0.001; SiO_4_, R=0.38, p=0.001; PO_4_^3-^, R=0.38, p=0.001) emerged as the main drivers (Fig. 2B) but others, such as salinity, fluorescence, and depth also showed significant association to the ordination of samples (Table S1). Temperature, salinity, and the trophic status have also been defined as key variables influencing the diversity and composition of AAP communities in the surface ocean [26, 27, 42]. A study based on the surface global ocean showed that AAP communities are subjected to strong selection processes even when there are small changes in the environmental conditions [28]. The Poseidon sampling covered four Longhurst provinces, each characterized by a unique combination of environmental variability (Fig. S5) that may promote the selection of specific taxa within provinces. Indeed, we found a separation of samples according to their Longhurst province (*envfit* analysis, R²=0.32, p<0.001, Fig. S6). This biogeographic classification has proven to be effective in explaining the spatial structure of various microbial communities [33, 52–55], including those of AAP bacteria ([28], this study). It is widely recognized that the environmental variability associated to depth is a major driver of changes in the composition of prokaryotic and eukaryotic communities [8–10, 56–59] but in contrast, the effects of that vertical component in the distribution of AAP communities have barely been explored. To illustrate the vertical variability in our dataset, we classified the samples by their position along the DCM profile (surface, above DCM, DMC and below DCM; Fig. 2A) rather than their depth, in order to better capture structure patterns linked to the DCM. We observed an ordination based on these layers (*envfit* analysis, R²=0.15, p<0.01), indicating that community structure is influenced by their position along the DCM profile. A few exceptions to this ordination correspond to samples from Station 10 (second DCM, at 100 m), Station 12 (below DCM, 170 m), Station 13 (below DCM, 210 m) and Station 14 (below DCM, 170 m), which were located in the oligotrophic gyre and characterized by deeper and weaker DCMs (Fig. 1B and Fig. 2B). A previous study based on *pufM* clone libraries in the Mediterranean Sea showed that populations above the DCM were more similar to each other than to those in or below the DCM [29]. In this study, we have significantly expanded the vertical resolution by considering different layers along the DCM structure and our results show that, besides the influence that can be attributed to the Longhurst provinces, the DCM structure influences the composition of AAP communities in the epipelagic zone.

### 3) Ca. *Luxescamonaceae* is the dominant AAP lineage in the euphotic ocean

We assigned the *pufM* gene sequences to specific taxonomic groups based on their placement in a phylogenetic tree (Fig. 3). Our classification used the phylogroups defined in Yutin et al. [42] complemented with the GTDB taxonomy, considering different orders of the classes Alphaproteobacteria (Sphingomonadales, Rhodobacterales, and Rhizobiales) and Gammaproteobacteria (Pseudomonadales and Burkholderiales). The 3,350 *pufM* amplicons were classified into 13 distinct taxonomic groups. Five of these groups –phylogroups A, B, C1, C2, D, and “Other Luxescamonaceae” (Fig. 3)– do not have cultured representatives and contain *pufM* sequences extracted from metagenomes, metagenome-assembled genomes (MAGs), and single amplified genomes (SAGs) from various oceanographic expeditions, including the *Tara* Oceans [10, 32, 60] and the Malaspina Expeditions [58, 61], and the Blanes Bay Microbial Observatory (BBMO) time series [49].

**Figure 3.**
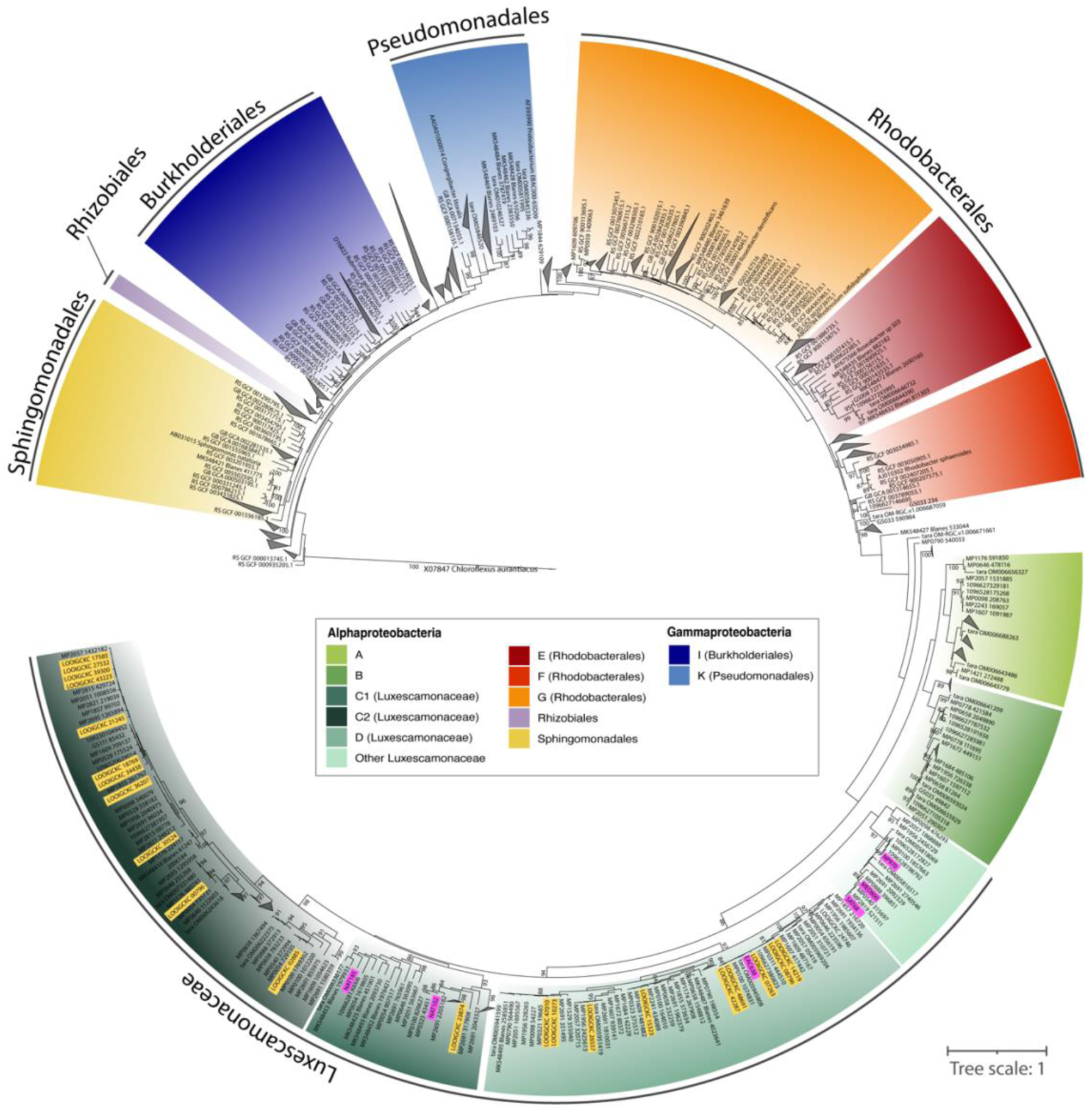
Phylogenetic tree of *pufM* sequences. Purple labels indicate *pufM* sequences related to *Candidatus* Luxescamonaceae from the metagenomic assembled genomes (MAGs) described in Graham et al. [31]. Yellow labels indicate *pufM* sequences related to *Ca.* Luxescamonaceae from single amplified genomes (SAGs) identified by Pachiadaki et al. [32]. Only bootstrap support values above 80 are displayed.

Phylogroup C2 consistently dominated along most stations and depths (mean relative abundance ± s.d. 37.68% ± 1.44), followed by phylogroup A (13.51% ± 7.80), phylogroup K within the Halieaceae family of the Pseudomonadales order (10.06% ± 6.89), phylogroup D (9.79% ± 6.13), phylogroup B (8.41% ± 10.83), and phylogroup G within the Rhodobacterales (6.76% ± 5.69) (Fig. 4, Table S2, Fig. S7). Other taxonomic groups exhibited mean relative abundances below 3%. Overall, the taxonomic composition of the AAP communities was strongly structured by the DCM profile rather than by depth (Fig. 4). Clear shifts in community composition occurred across regions with contrasting oceanographic conditions and different DCM depths. For instance, in Stations 12, 13, and 14, located within the oligotrophic gyre where the DCM reached ∼150 m, community changes at the DCM were marked by increases in a few ASVs from phylogroup K (Pseudomonadales) and Rhizobiales. In contrast, stations in the north of the equator (Stations 19–22, Fig. 4), with shallower and more intense chlorophyll peaks (>1 mg·m⁻³), exhibited shifts in taxonomic composition, with an increase in the abundance of phylogroups A and B, “Other Luxescamonaceae”, and a decrease in the relative abundance of phylogroup G (Rhodobacterales). In addition, AAP assemblages varied across oceanic regions. For example, Station 2, influenced by Southern Ocean conditions, harbored several taxa from the Sphingomonadales and phylogroup E (Rhodobacterales), which were nearly absent elsewhere (Fig. 4).

**Figure 4.**
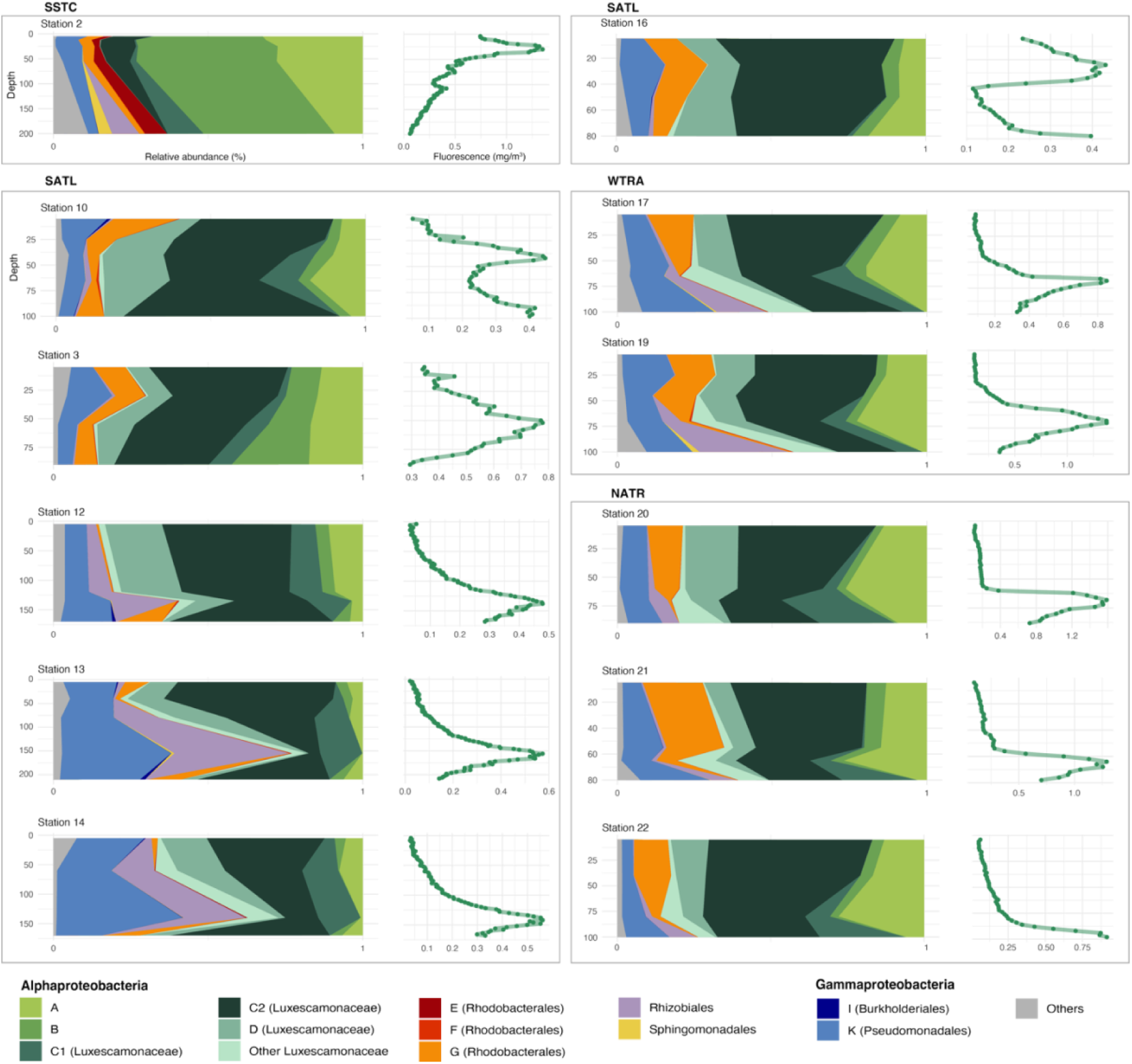
Taxonomic composition of AAP bacterial communities (left of each panel) and depth fluorescence variation (right of each panel) across depths in twelve stations along the Poseidon Expedition transect (only stations with more than three sampling points are shown). Stations from different Longhurst provinces are delimited within different boxes. SSTC, South Subtropical Convergence zone; SATL, South Atlantic gyre, WTRA, Western tropical Atlantic, and NATR, North Atlantic Tropical gyre.

Contrary to previous assumptions, our results challenge the prevailing view that Pseudomonadales (phylogroup K) and Rhodobacterales (particularly phylogroup G) dominate AAP assemblages in the open ocean [26, 28, 29, 46]. The taxonomic composition observed here is similar to that reported in Yutin et al., [42], which was based solely on metagenomic data. In fact, our choice of primers follows Gazulla et al. [30], in which we demonstrated that traditionally employed primers have multiple mismatches for phylogroups A, B, C, and D, leading to their underestimation. By using primers with higher phylogenetic coverage, we revealed that indeed these groups constitute a substantial fraction of AAP communities along both the vertical and horizontal continuum. Interestingly, little ecological or physiological information exists for these groups, as no representatives isolates are currently available. The phylogenetic analyses show that phylogroups C and D cluster with sequences from *Ca*. Luxescamonaceae (Bootstrap Support=0.86, Fig. 3), a family with the genomic potential for carbon fixation (containing ribulose-1,5-bisphosphate carboxylase (RuBisCO) form IC/D, [31]). First reported from the *Tara* Oceans metagenomic dataset [31], the clade *Ca*. Luxescamonaceae was later described in single amplified genomes (SAGs) from the same expedition [32]. These studies place it within the Alphaproteobacteria, and a recent study defines them as the LUX cluster within the *Roseobacter* group [62]. In addition, our phylogenetic tree shows that this cluster could be further divided into four subclusters: two robust sub-clusters (BS > 95), phylogroup D and part of phylogroup C (cluster C2), and another two sub-clusters with lower confidence level that represent phylogroup C (cluster C1, BS=0.35) and other sequences that could not be assigned to any of these phylogroups, and that we named “Other Luxescamonaceae” (BS=0.68, Figure 3A). Previous studies analyzing the phylogeny of *pufM* sequences show very similar phylogenetic clustering for phylogroups C and D [28, 50]. In our study, a total of 1,579 ASVs (47.13% of total number of ASVs) were assigned to the clusters within the *Ca.* Luxescamonaceae family, representing more than half of the total AAP relative abundance (54.86% ± 14.53, Table S2). Among them, we found contrasting vertical distribution patterns. Luxescamonaceae C2 was more abundant at the surface and decreased progressively through the DCM profile, whereas Luxescamonaceae C1 peaked below the DCM (Fig. 5A). Clade C1 thrived in deeper, nutrient-rich and colder waters while clades C2 and D correlate with the higher salinity and warmer temperatures found in surface waters of the Atlantic Ocean (Fig. 5B). Based on SAGs and MAGs recovered from tropical to temperate waters across major oceans and adjacent seas, the LUX cluster comprises genomes showing significant differences in genome features such as genome size, G+C content, and coding density [62]. These genomic differences, together with the observed variation in ecological niches observed in our study, suggest that *Ca.* Luxescamonaceae is not a single cohesive ecological group but rather contains various ecotypes adapted to the distinct conditions within the euphotic ocean. Although not experimentally confirmed, their genomic repertoire suggest that some members of *Ca.* Luxescamonaceae may be capable of carbon fixation, implying a mix of phototrophy and photoheterotrophy [31]. While phylogroups C and D are known to be related to *Ca.* Luxescamonaceae, much less is known about phylogroups A and B. Initially identified in oligotrophic regions of the Atlantic and Pacific oceans [42], they were recently found to dominate AAP communities in the Mediterranean Sea, particularly during early spring and winter [63]. In our study, phylogroup A consistently peaked at the DCM and showed a positive correlation with Chl *a* concentration and fluorescence (Pearson corr. with Chl *a* R=0.56, p<0.001; fluorescence R=0.57, p<0.001, Fig. 3C). Phylogroup B was generally scarce across the dataset, except at Station 2, where it accounted for approximately 40% of the community from the surface down to 200 m depth. This station, located in the SSTC region and highly influenced by its proximity to the Southern Ocean, had a clearly different AAP community, as revealed by ordination analyses (Fig. S6). These observations suggest that phylogroup B may be better adapted to high-latitude, cold-water environments.

**Figure 5.**
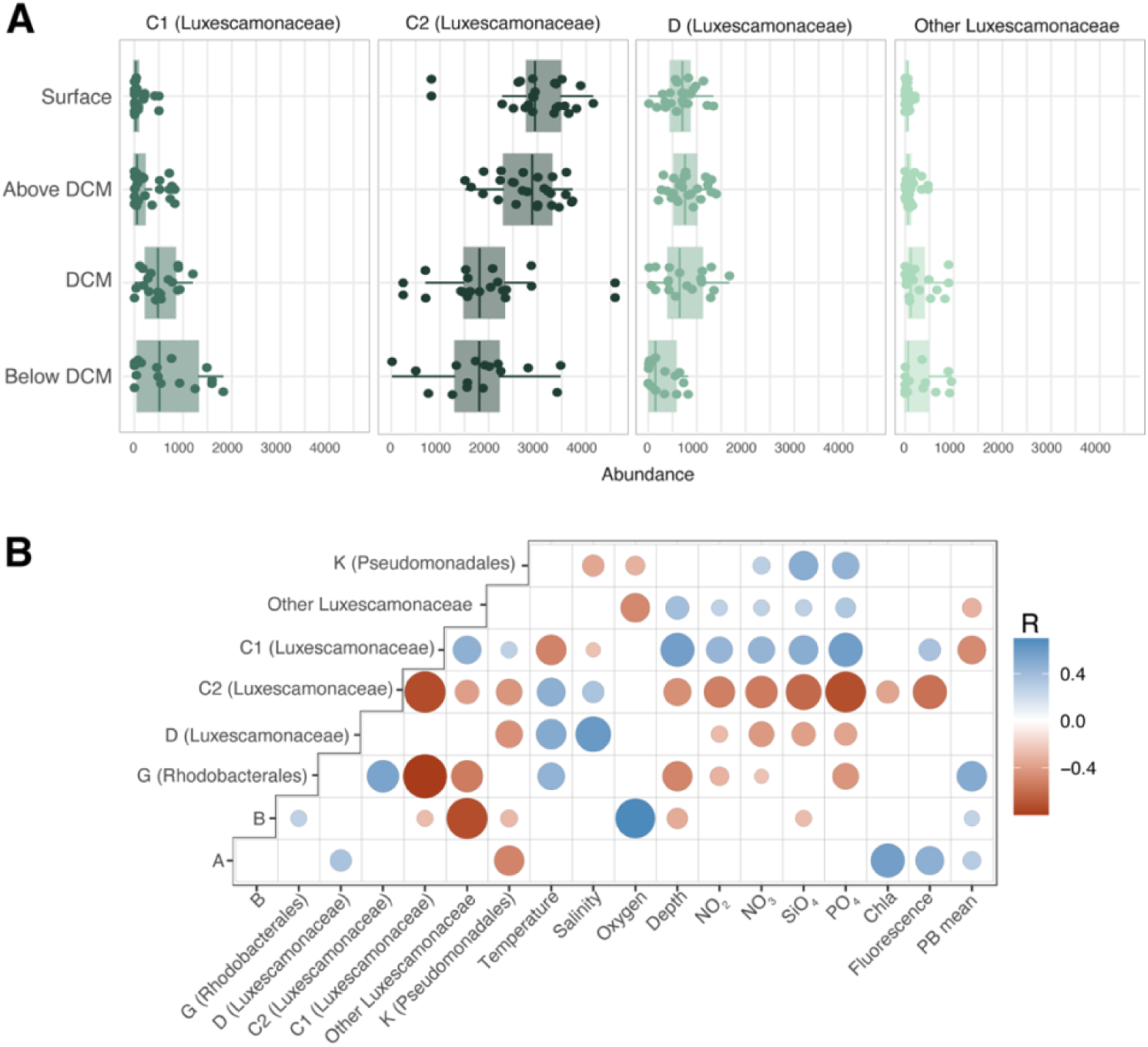
**A)** Distribution of abundance of the four sub-clusters classified as *Ca.* Luxescamonaceae, along the deep chlorophyll maximum profile. Read counts represent values after rarefaction to 6,576 reads. **B)** Pearson correlation coefficients between the relative abundance of the most prevalent taxonomic groups (mean relative abundance above 3%) and environmental variables. The plot only displays statistically significant (p<0.05) correlations (R) after Bonferroni correction. Chla: chlorophyll *a* concentration, PB mean: Heterotrophic bacterial production

Although typically studied collectively as a cohesive microbial guild, AAP bacteria comprise very diverse metabolisms and life strategies, ranging from generalists to specialists that thrive on different ranges of carbon sources [64], and with distinct habitat preferences [27, 49, 63]. By integrating vertical patterns across the subtropical and tropical ocean, this study provides new insight into the ecological preferences of AAPs and, for the first time, describes the ecological distribution of the abundant Luxescamonaceae family within these communities. It remains unknown whether the *pufM* gene sequences recovered in our study originate from genomes with carbon fixation potential, or if so, whether these genes are functionally active. If they are, given the high abundance of Luxescamonaceae throughout the Atlantic Ocean, the ecological role traditionally attributed to AAP bacteria would necessitate a reevaluation.

## Conclusions

This study shows for the first time that AAP bacterial diversity and community structure closely follow the deep chlorophyll maximum (DCM), with richness peaking at or just above the DCM and correlating with chlorophyll *a* concentration. Temperature, oxygen, and nutrient concentrations are the primary environmental drivers shaping AAP communities, with salinity and vertical position within the DCM also contributing. Biogeographic patterns are further structured by Longhurst provinces, highlighting the role of environmental selection. Our results reveal that previously underestimated uncultured phylogroups (A, B, C, and D) constitute a substantial fraction of AAP communities, likely overlooked due to primer biases. Remarkably, *Candidatus* Luxescamonaceae dominates the Atlantic Ocean euphotic zone, comprising multiple subclusters with distinct vertical distributions, suggesting different ecotypes adapted to varying environmental conditions. Overall, these findings underscore the ecological diversity and environmental specialization of AAP bacteria, while highlighting important knowledge gaps that could affect our understanding of their contribution to marine phototrophic processes.

## Acknowledgments

We would like to thank the entire crew of the R/V Sarmiento de Gamboa for their invaluable support during the POSEIDON Expedition. We would like to thank Paula Sabaté and Oriol Puig for help with DNA extractions. We also extend our gratitude to the MARBITS bioinformatics platform at the Institut de Ciències del Mar, as well as the Picasso Supercomputer at the Supercomputing and Bioinformatics Center of the University of Málaga, and their staff, for providing the necessary computing resources and technical assistance.

## Author contributions

CRG, IF, JMG and OS conceived the study; IF, JMA, JMG, and OS acquired funding; IF, VB, CMV, AGV, JEP, EFN, and JMA collected data and samples along the cruise; CRG, VB, CMV, and JEP processed the samples. CRG analyzed the data and wrote the first draft of the manuscript with significant contributions from IF, JMG, and OS. All authors reviewed the manuscript.

