## Supplementary material for "Diversity and community structure of aerobic anoxygenic phototrophic bacteria are shaped by the deep chlorophyll maximum": supp_material: Gazulla2025_AAPsPoseidon_bioRxiv.docx

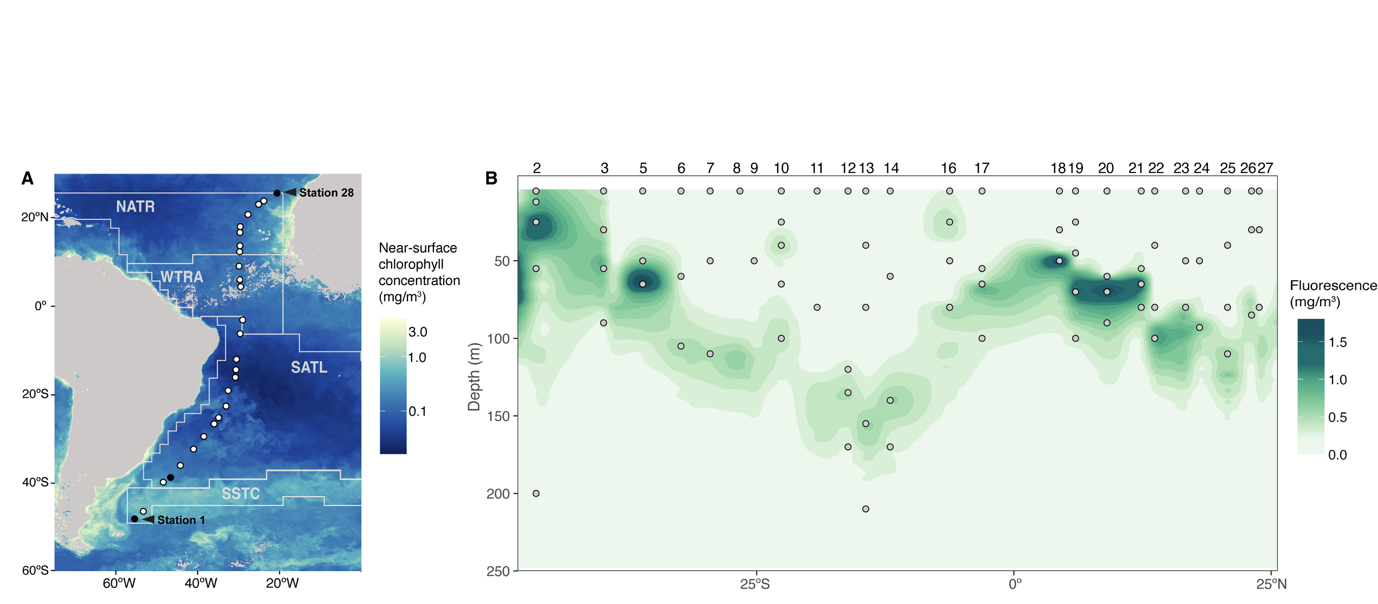

**Figure 1. A)** Near-surface chlorophyll *a* concentrations and station positions along the Poseidon Expedition transect in the South and Central Atlantic (March 2019). Colors indicate chlorophyll *a* (mg·m⁻³) from SNPP-VIIRS data (<https://oceancolor.gsfc.nasa.gov/l3/>. The transect covers four Longhurst provinces: SSTC, South Subtropical Convergence zone; SATL, South Atlantic gyre, WTRA, Western tropical Atlantic, and NATR, North Atlantic Tropical gyre. **B)** Sectional distribution of *in situ* chlorophyll fluorescence measured by the CTD along the transect. Numbers above the panel indicate station number. Dots indicate the sampling points. Note that station 15 does not exist. Figure adapted from Gazulla et al. [20].

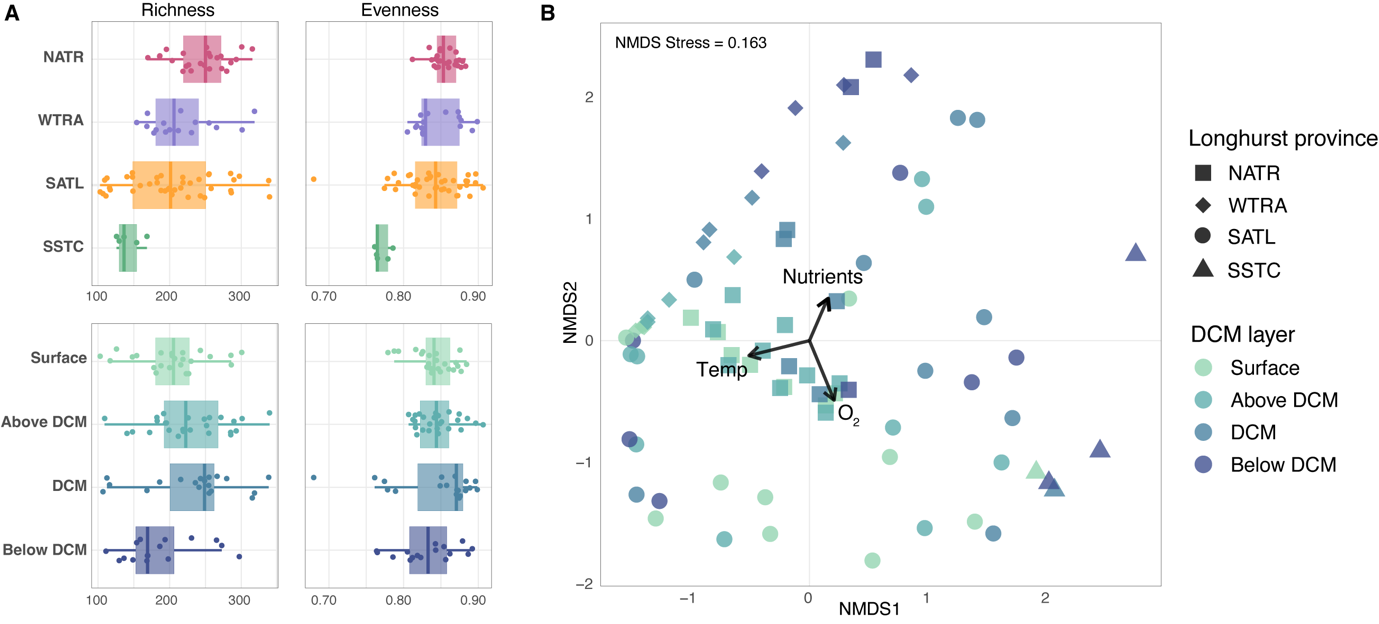

**Figure 2. A)** Alpha diversity measured across latitude (horizontal scale, top panels) and vertically through the Deep Chlorophyll Maximum (DCM) structure (bottom panels). Richness was measured using the Chao1 index. **B)** Non-metrical multidimensional (nMDS) plot based on the Bray-Curtis dissimilarities of AAP community composition. Samples from the distinct DCM layers and Longhurst provinces are denoted by different colors and shapes, respectively. The *envfit* analysis identified temperature (Temp), oxygen (O_2_) and nutrients as the three variables that explained the largest fraction of community variance. SSTC, South Subtropical Convergence zone; SATL, South Atlantic gyre, WTRA, Western tropical Atlantic, and NATR, North Atlantic Tropical gyre.

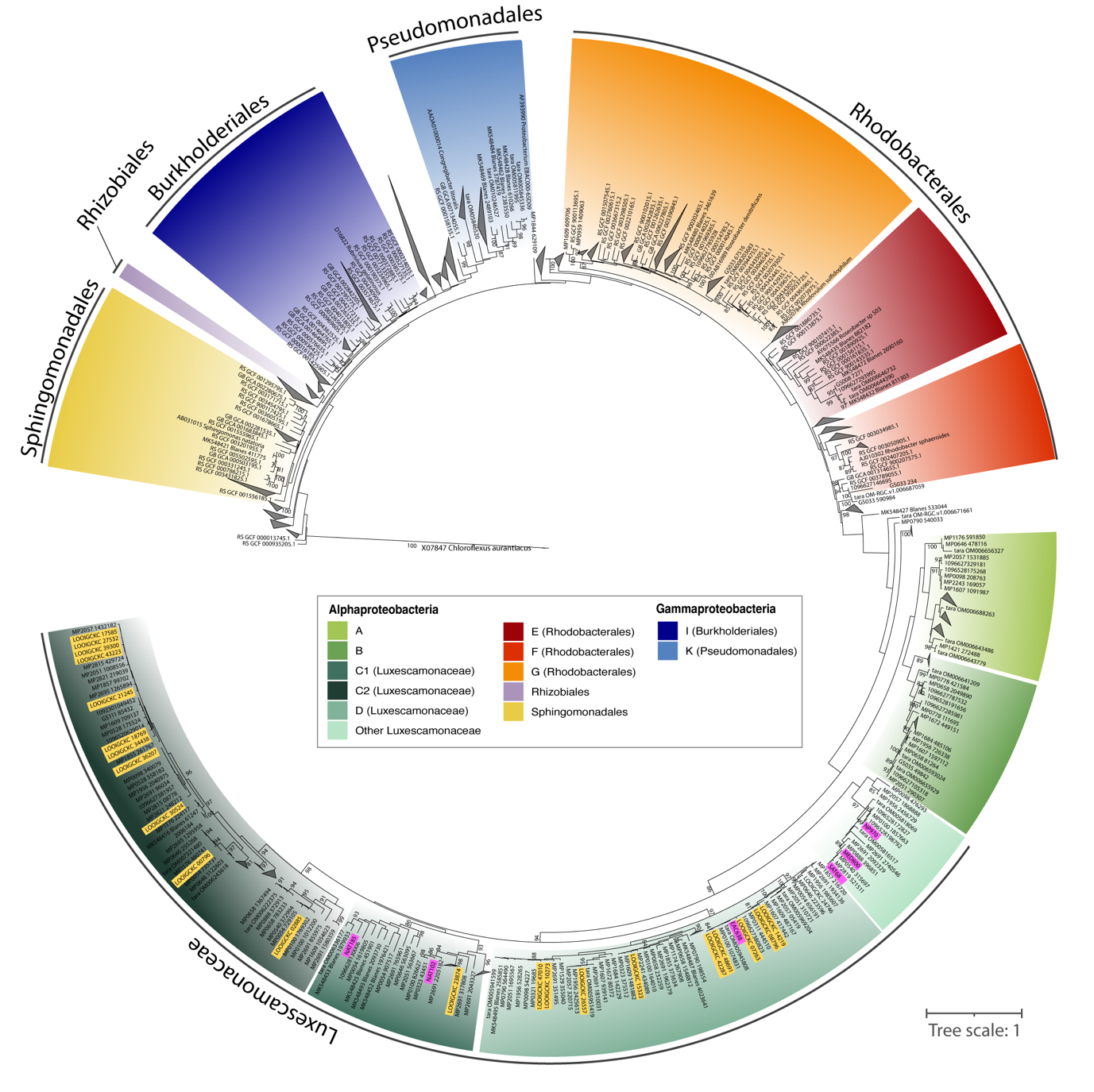

**Figure 3.** Phylogenetic tree of *pufM* sequences. Purple labels indicate *pufM* sequences related to *Candidatus* Luxescamonaceae from the metagenomic assembled genomes (MAGs) described in Graham et al. [31]. Yellow labels indicate *pufM* sequences related to *Ca.* Luxescamonaceae from single amplified genomes (SAGs) identified by Pachiadaki et al. [32]. Only bootstrap support values above 80 are displayed.

Phylogroup C2 consistently dominated along most stations and depths (mean relative abundance ± s.d. 37.68% ± 1.44), followed by phylogroup A (13.51% ± 7.80), phylogroup K within the Halieaceae family of the Pseudomonadales order (10.06% ± 6.89), phylogroup D (9.79% ± 6.13), phylogroup B (8.41% ± 10.83), and phylogroup G within the Rhodobacterales (6.76% ± 5.69) (Fig. 4, Table S2, Fig. S7). Other taxonomic groups exhibited mean relative abundances below 3%. Overall, the taxonomic composition of the AAP communities was strongly structured by the DCM profile rather than by depth (Fig. 4). Clear shifts in community composition occurred across regions with contrasting oceanographic conditions and different DCM depths. For instance, in Stations 12, 13, and 14, located within the oligotrophic gyre where the DCM reached ~150 m, community changes at the DCM were marked by increases in a few ASVs from phylogroup K (Pseudomonadales) and Rhizobiales. In contrast, stations in the north of the equator (Stations 19–22, Fig. 4), with shallower and more intense chlorophyll peaks (>1 mg·m⁻³), exhibited shifts in taxonomic composition, with an increase in the abundance of phylogroups A and B, “Other Luxescamonaceae”, and a decrease in the relative abundance of phylogroup G (Rhodobacterales). In addition, AAP assemblages varied across oceanic regions. For example, Station 2, influenced by Southern Ocean conditions, harbored several taxa from the Sphingomonadales and phylogroup E (Rhodobacterales), which were nearly absent elsewhere (Fig. 4).

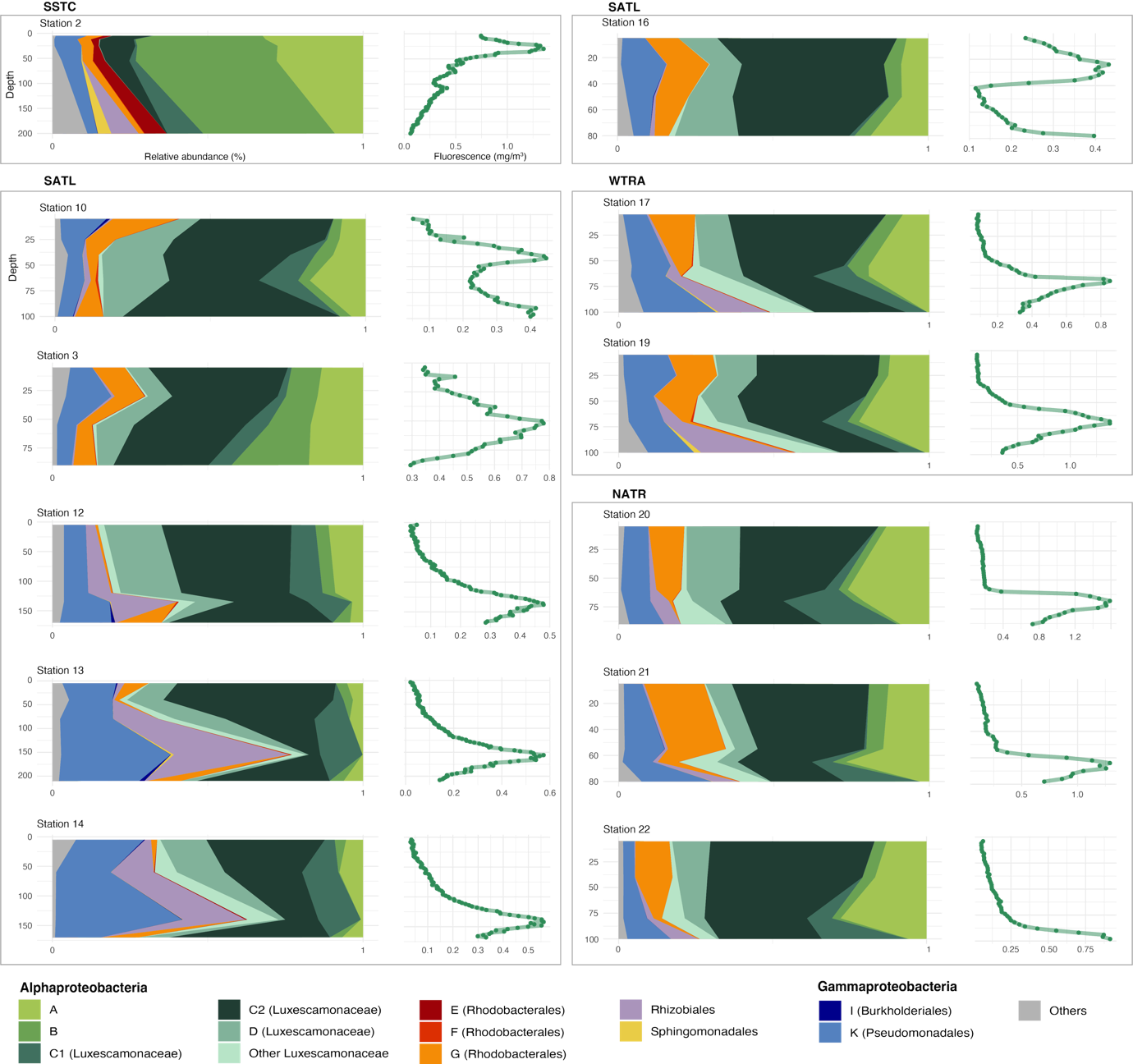

**Figure 4.** Taxonomic composition of AAP bacterial communities (left of each panel) and depth fluorescence variation (right of each panel) across depths in twelve stations along the Poseidon Expedition transect (only stations with more than three sampling points are shown). Stations from different Longhurst provinces are delimited within different boxes. SSTC, South Subtropical Convergence zone; SATL, South Atlantic gyre, WTRA, Western tropical Atlantic, and NATR, North Atlantic Tropical gyre.

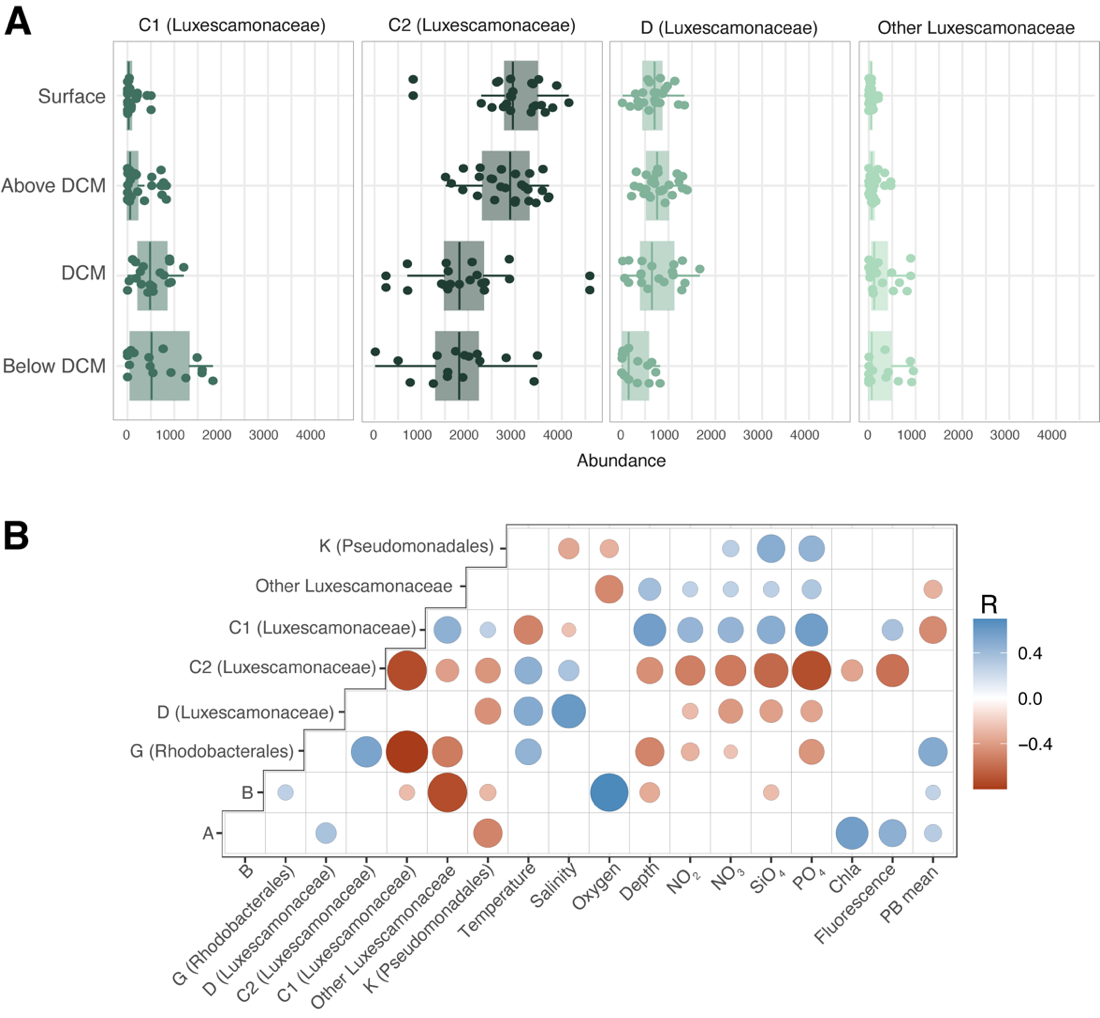

**Figure 5. A)** Distribution of abundance of the four sub-clusters classified as *Ca.* Luxescamonaceae, along the deep chlorophyll maximum profile. Read counts represent values after rarefaction to 6,576 reads. **B)** Pearson correlation coefficients between the relative abundance of the most prevalent taxonomic groups (mean relative abundance above 3%) and environmental variables. The plot only displays statistically significant (p<0.05) correlations (R) after Bonferroni correction. Chla: chlorophyll *a* concentration, PB mean: Heterotrophic bacterial production

**Conflicts of interest**

The authors have no conflicts of interest to declare.

**Funding**

This work was supported by the ECLIPSE grant (PID2019-110128RB-I00/AEI/10.13039/501100011033) to IF, the MICOLOR grant to JMG (PID2021-125469NB-C31) and OS (PID2021-125469NB-C32), and the POSEIDON grant (CTM2017-84735-R) to JMA, all funded by the Agencia Estatal de Investigación of the Spanish Ministry of Science, Innovation and Universities (MCIN/AEI/10.13039/501100011033/FEDER, UE). Authors affiliated with the Institut de Ciències del Mar received institutional support through the ‘Severo Ochoa Centre of Excellence’ accreditation (CEX2024-001494-S). CRG was supported by a PIF fellowship from the Universitat Autònoma de Barcelona.

**Author contributions**

CRG, IF, JMG and OS conceived the study; IF, JMA, JMG, and OS acquired funding; IF, VB, CMV, AGV, JEP, EFN, and JMA collected data and samples along the cruise; CRG, VB, CMV, and JEP processed the samples. CRG analyzed the data and wrote the first draft of the manuscript with significant contributions from IF, JMG, and OS. All authors reviewed the manuscript.
